# Intrinsic representational drift from predictive excitatory-inhibitory plasticity

**DOI:** 10.64898/2026.06.17.733055

**Authors:** Toshitake Asabuki, Claudia Clopath

## Abstract

Neural representations drift gradually over time even under stable environmental conditions, but the synaptic mechanisms driving this drift remain unclear. Here we show that representational drift can arise intrinsically from predictive synaptic plasticity in spiking excitatory-inhibitory networks. During repeated exposure to unchanged input patterns, individual neurons gradually changed their preferred pattern while ensemble-level coding remained stable. These changes in preference were preceded by a weakening of the net excitatory-inhibitory drive supporting the current preference relative to competing patterns, and this relative drive predicted changes on the following trial. Extending the model to hippocampal place coding reproduced experience-dependent tuning-curve drift in CA1, including the dissociation between elapsed time and intervening exposure. At the population level, drift was expressed as coordinated rotation and translation of neural state space. Thus, representational drift can emerge as an intrinsic consequence of predictive E/I plasticity that maintains balanced, selective population codes.

## Introduction

The stability of memories and behaviors is often thought to rely on the stability of the underlying neural representations. Hippocampal place cells, for example, provide an apparently robust map of familiar environments by firing at specific locations (O’Keefe & Dostrovsky, 1971). Yet chronic recordings over days to weeks have revealed that these representations are not static. In hippocampal CA1, many neurons change their preferred firing locations or cease to be active across days, even while the population continues to encode spatial information reliably (Ziv et al., 2013; Rubin et al., 2015; Hainmueller & Bartos, 2018). Similar representational drift has been observed in parietal cortex, visual areas, and olfactory cortex (Driscoll et al., 2017; Deitch et al., 2021; Marks & Goard, 2021; Schoonover et al., 2021), suggesting that ongoing representational change is a general feature of cortical circuits rather than a hippocampal specialization. At the population level, this drift has recently been characterized as a gradual rotation and translation of the neural state space, with the relational geometry of the code partly preserved even as individual tuning profiles change (Sylte et al., 2025). These observations have motivated broader theoretical accounts of how neural representations can remain informative despite long-term variability and partial instability (Clopath et al., 2017; Gonzalez et al., 2019; Driscoll et al., 2022; Rule & O’Leary, 2022; Delamare et al., 2024).

Recent experiments have further shown that representational drift is not simply random degradation. In CA1, the identities and tuning properties of individual neurons can change over days to weeks, while population-level information is partially preserved (Rubin et al., 2015; Sadeh & Clopath, 2022). Moreover, time and experience affect different aspects of representational stability. Ensemble firing rates and spatial tuning curves show distinct dependencies on elapsed time and the number of intervening experiences, and active experience can drive representational change even within a day (Geva et al., 2023; Khatib et al., 2023). Related studies of engram dynamics further show that neuronal ensembles are not fixed entities. Instead, ensemble membership is shaped by experience history and cellular state, which bias memory allocation and subsequent ensemble remodeling over time (Cai et al., 2016; Cho et al., 2021; Silva et al., 2009; Zhou et al., 2009; Rashid et al., 2016; Rogerson et al., 2014; Pignatelli et al., 2019; Josselyn & Tonegawa, 2020; Mau et al., 2020; Delamare et al., 2024). These findings suggest that representational drift reflects an interaction between ongoing circuit changes and experience-dependent plasticity. However, the synaptic mechanisms linking experience, plasticity, and individual neuronal changes remain unclear. In particular, it remains unknown what determines when and why individual neurons change their representational preference.

Several mechanisms could contribute to drift, including structural synaptic turnover (Attardo et al., 2015), ongoing Hebbian or homeostatic plasticity (Vogels et al., 2011; Qin et al., 2023), intrinsic excitability fluctuations (Delamare et al., 2024), and experience- or behavior-dependent changes in neural activity (Rule et al., 2019; Sadeh & Clopath, 2022; Khatib et al., 2023). Computational models have shown that externally imposed synaptic noise or weight perturbations can produce gradual changes in neural activity patterns (Rule et al., 2019). Other models invoke Hebbian competition or attractor dynamics in which stochastic fluctuations destabilize individual neurons while the network as a whole remains informative (Kossio et al., 2021; Qin et al., 2023; Ratzon et al., 2024). However, in many of these approaches, the changes that drive drift are treated as external perturbations or phenomenological noise. It therefore remains unclear what synaptic process determines when and why individual neurons change their preferred representation, and how such single-neuron changes can coexist with stable population-level coding.

Here we address this problem using a recurrent spiking excitatory-inhibitory network trained with a local predictive learning rule, in which excitatory synapses predict postsynaptic activity and inhibitory synapses track excitatory drive to maintain predictive E/I balance. Repeated exposure to the same input patterns was sufficient to produce representational drift. Trial-resolved drive analysis showed that changes in preference were preceded by a weakening of the net drive supporting the current preference, partly through progressive strengthening of inhibition onto the active assembly. This relative drive predicted changes on the following trial, linking ongoing E/I rebalancing to individual preference changes. Extending the same framework to hippocampal place coding reproduced recently reported experimental findings, including separable time- and experience-dependent stability (Geva et al., 2023) and coordinated rotation and translation of population state space (Sylte et al., 2025). These results provide a synaptic mechanism linking local predictive plasticity to single-neuron instability and population-level stability.

## Results

### Local predictive plasticity generates pattern-selective assemblies that drift over time

We asked whether local predictive plasticity could account for both the emergence of pattern-selective assemblies and their gradual reorganization over time. We used a recurrent network driven by three structured input patterns (Fig. 1A). Feedforward excitatory, recurrent excitatory, and recurrent inhibitory synapses were updated by a local predictive learning rule (Fig. 1B; Asabuki & Fukai, 2025; Asabuki & Clopath, 2025). Excitatory synapses learned to predict postsynaptic activity from feedforward or recurrent excitatory drive, whereas inhibitory synapses tracked excitatory drive and stabilized the resulting activity.

**Figure 1.**
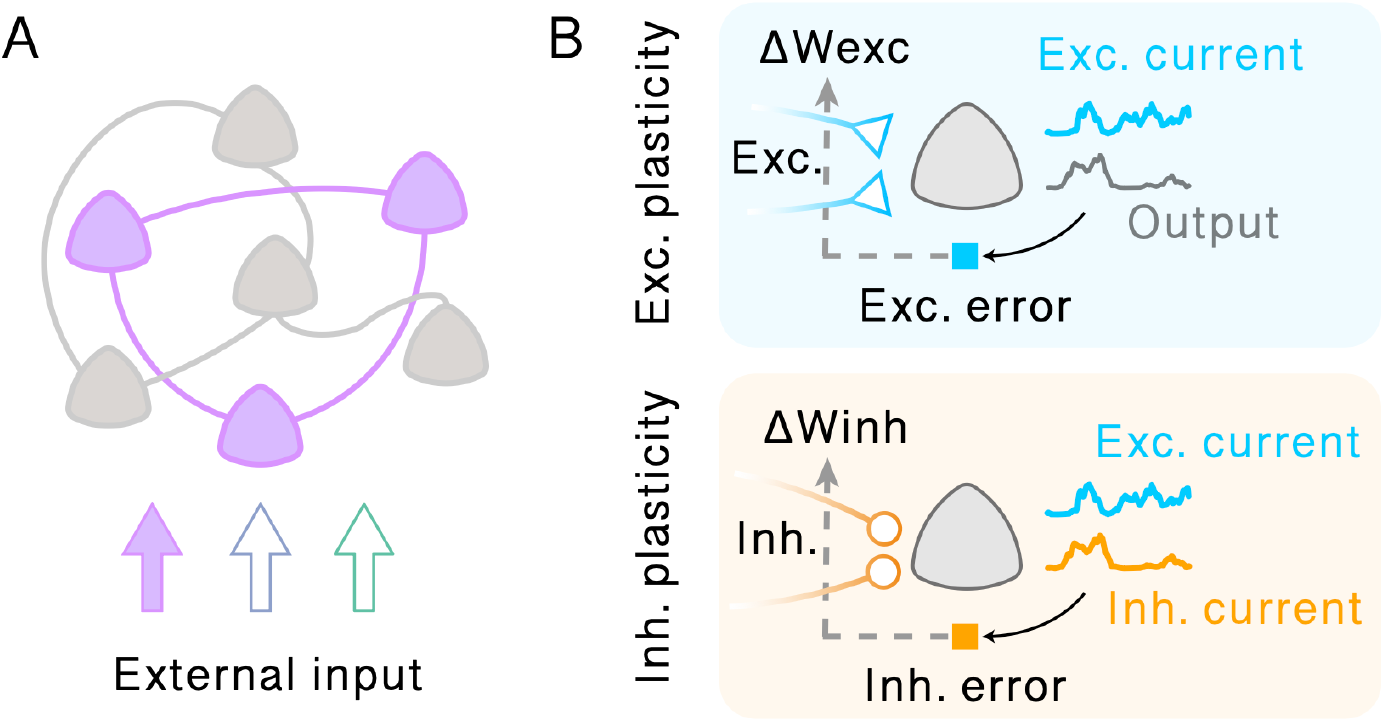
Predictive learning rule in spiking recurrent networks. (A) Schematic of the spiking recurrent network model. (B) A schematic of the plasticity rules. Excitatory (blue; feedforward and recurrent) and inhibitory (orange; recurrent) synapses projecting to a postsynaptic neuron (triangle) obey different plasticity rules. For excitatory synapses, errors between excitatory drive and the output of the cell provide feedback to the synapses (dashed arrow) and modulate plasticity (blue square; exc. error). All excitatory connections seek to minimize these errors. For inhibitory synapses, the error between excitatory and inhibitory drive must be minimized to maintain excitation–inhibition balance (orange square; inh. error).

The network developed pattern-selective assemblies during training (Fig. 2A; Day 1). We then tracked how this population structure evolved during continued learning under the same input statistics. Although the same input patterns were repeatedly presented, the membership of each assembly progressively reorganized. Indeed, when activity on Day 10 was visualized using the neuron ordering defined on Day 1, the original assembly structure was disrupted (Fig. 2A; Day 10), suggesting that neurons no longer belonged to the same pattern-selective assemblies as in the reference state. To determine whether this reflected a loss of assembly organization or a reorganization of assembly membership, we examined pairwise correlation matrices using different sorting orders. Clear block-like structure was observed when activity from each day was sorted by the preferences defined on that same day, but was weakened when sorted by preferences from a different day (Fig. 2B). Thus, assemblies remained organized at each time point, but the neurons composing each assembly changed across days.

**Figure 2.**
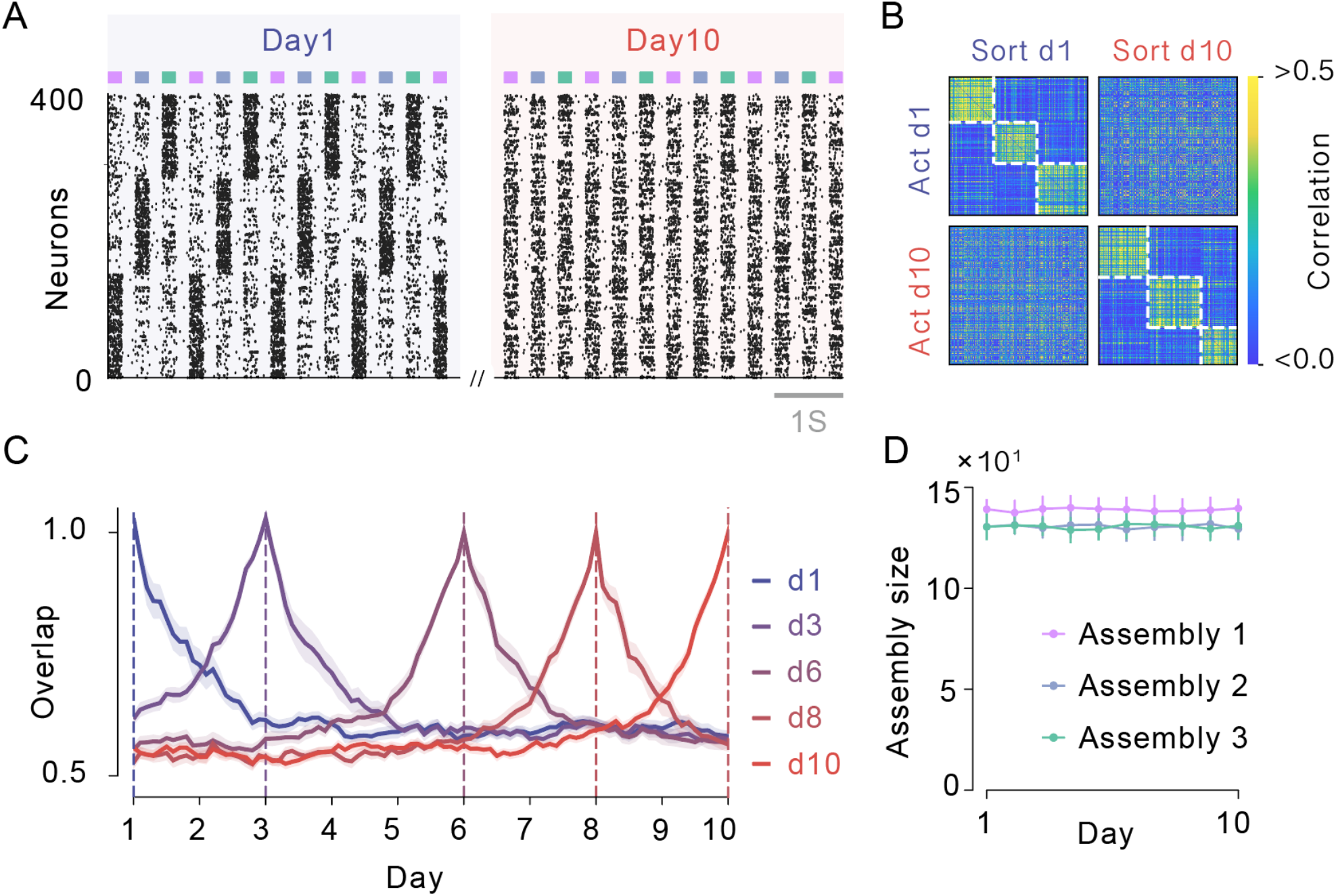
Pattern-selective assemblies drift while population structure remains organized. (A) Evoked population activity on Day 1 and Day 10, with neurons sorted by their preferred pattern on Day 1. The Day 1 ordering reveals clear assemblies, whereas the same ordering shows disrupted structure on Day 10. (B) Pairwise correlation matrices of evoked activity, with rows indicating the activity day and columns indicating the sorting day. Block-like structure is clear only when activity is sorted by preferences from the same day. (C) Assembly overlap across the drift period, computed for multiple reference snapshots. (D) Assembly sizes remained approximately stable across the drift period.

We quantified this turnover by measuring assembly overlap across the drift period. Assembly overlap decayed gradually with temporal distance from the reference snapshot, and similar decay profiles were observed when different snapshots were used as references (Fig. 2C). This indicates that drift proceeded continuously rather than reflecting a transient early reorganization. Despite this turnover, assembly sizes remained approximately stable across the drift period (Fig. 2D), suggesting that drift reflected reassignment of neurons between assemblies rather than systematic expansion or collapse of individual assemblies. Consistently, pattern identity remained perfectly decodable from population activity throughout the drift period (Supplementary Fig. 1).

Together, these results show that local predictive plasticity generates drifting but organized assemblies. Individual neurons gradually changed their assembly membership, yet the population retained a structured assembly organization throughout.

### E/I rebalancing drives single-neuron preference change

We next sought to identify the synaptic changes that precede preference changes in individual neurons. In the drifting assemblies described above, some neurons lost their association with one pattern-selective assembly and became reassigned to another assembly (Fig. 3A). Because the underlying drift was continuous, we operationally defined stable preference changes as changes in preferred pattern that persisted across consecutive trials. We then aligned these transitions to the time of reassignment and examined how pattern-conditioned synaptic drive changed around them.

**Figure 3.**
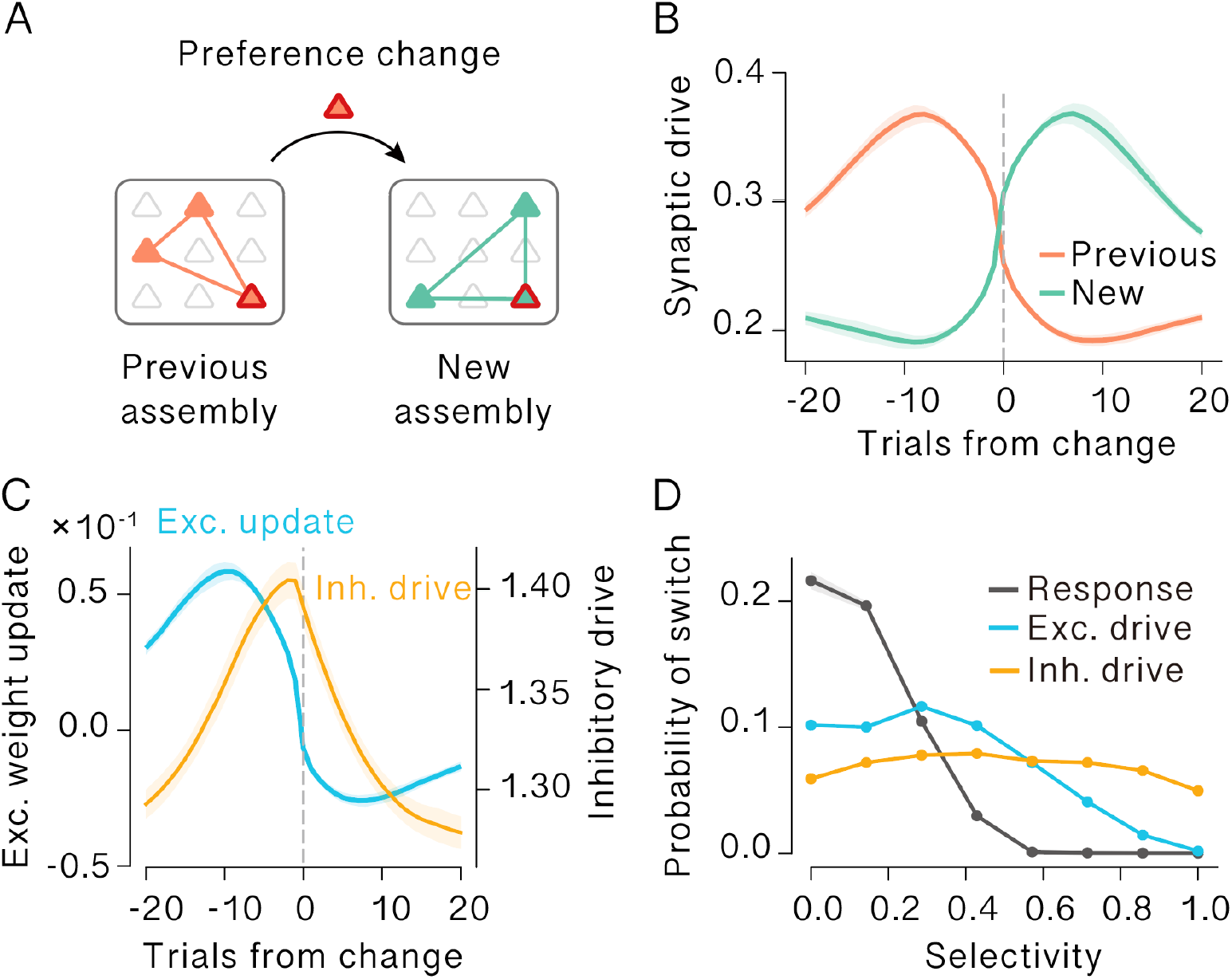
E/I rebalancing precedes and predicts single-neuron preference change. (A) Schematic of assembly change. A neuron initially assigned to the orange assembly is reassigned to the green assembly. (B) Transition-triggered average of pattern-conditioned net E/I drive (total excitatory minus inhibitory drive) onto neurons undergoing a preference change. Drive associated with the previously preferred pattern was dominant before the change but dropped sharply at reassignment, whereas drive associated with the newly preferred pattern rose before reassignment and became dominant afterward. (C) Excitatory weight update and inhibitory drive associated with the previously preferred pattern around change. The excitatory update (blue) decreased sharply before the change and became negative after reassignment. This decrease coincided with a transient increase in inhibitory drive (orange). (D) Probability of change on the next trial as a function of current selectivity. Selectivity was defined as the difference between the largest and second-largest pattern-conditioned values for each neuron. Higher response selectivity (black) and excitatory-drive selectivity (blue) predicted lower change probability, whereas inhibitory-drive selectivity (orange) showed little relationship with future change. In B and C, shaded regions indicate s.e.m. across 20 change events.

Neurons undergoing a preference change showed a stereotyped transition in net E/I drive. The net drive associated with the previously preferred pattern was dominant before the change but declined and dropped sharply around reassignment. Concurrently, the net drive associated with the newly preferred pattern began rising before the change and became dominant afterward (Fig. 3B). The preference change was therefore not an isolated abrupt event, but reflected a competition between the two assembly representations that resolved in favor of the new pattern.

To identify the synaptic mechanism underlying this net drive transition, we examined the excitatory weight update associated with the previously preferred pattern. Before the change, the excitatory update was positive, consistent with continued strengthening of synapses supporting the current assembly. Shortly before reassignment, however, this update decreased sharply and turned negative after the change (Fig. 3C). This reversal coincided with a transient increase in inhibitory drive. Because the excitatory update depends on the mismatch between firing activity and the excitatory prediction, the inhibition-driven suppression of firing reduces this mismatch below zero, thereby reversing the excitatory learning signal and weakening synapses that previously supported the current assembly.

We then asked whether the strength of a neuron’s current pattern preference predicted future change. For each neuron and trial, we defined selectivity as the difference between the largest and second-largest pattern-conditioned values. Response selectivity was computed from firing responses, excitatory-drive selectivity from excitatory synaptic drive, and inhibitory-drive selectivity from inhibitory drive. The probability of change on the next trial decreased strongly with response selectivity (Fig. 3D). A similar relationship was observed for excitatory-drive selectivity, whereas inhibitory-drive selectivity showed little relationship with future change. Thus, neurons were most likely to change when excitatory-drive support for the current preferred pattern had already weakened.

Together, these results indicate that single-neuron assembly change is driven by coordinated E/I rebalancing. A transient increase in inhibition suppresses firing, reverses the excitatory learning signal, and weakens synaptic support for the previous assembly. This process weakens synaptic support for the current preferred pattern and increases the probability of reassignment to another assembly.

### The model reproduces experience-dependent hippocampal representational drift

We next asked whether the same predictive plasticity framework could reproduce the time- and experience-dependent components of hippocampal representational drift reported by Geva et al. (2023). In that study, CA1 activity was recorded longitudinally while mice experienced two familiar environments at different frequencies, allowing elapsed time and intervening experience to be dissociated. We therefore simulated two linear-track environments with different exposure schedules: Env A was visited every 2 days, whereas Env B was visited every 4 days (Fig. 4A). During each environmental exposure, spatially tuned input neurons drove the recurrent network along the linear track. After learning, the model produced place-cell-like tuning curves that gradually changed across days (Fig. 4B).

**Figure 4.**
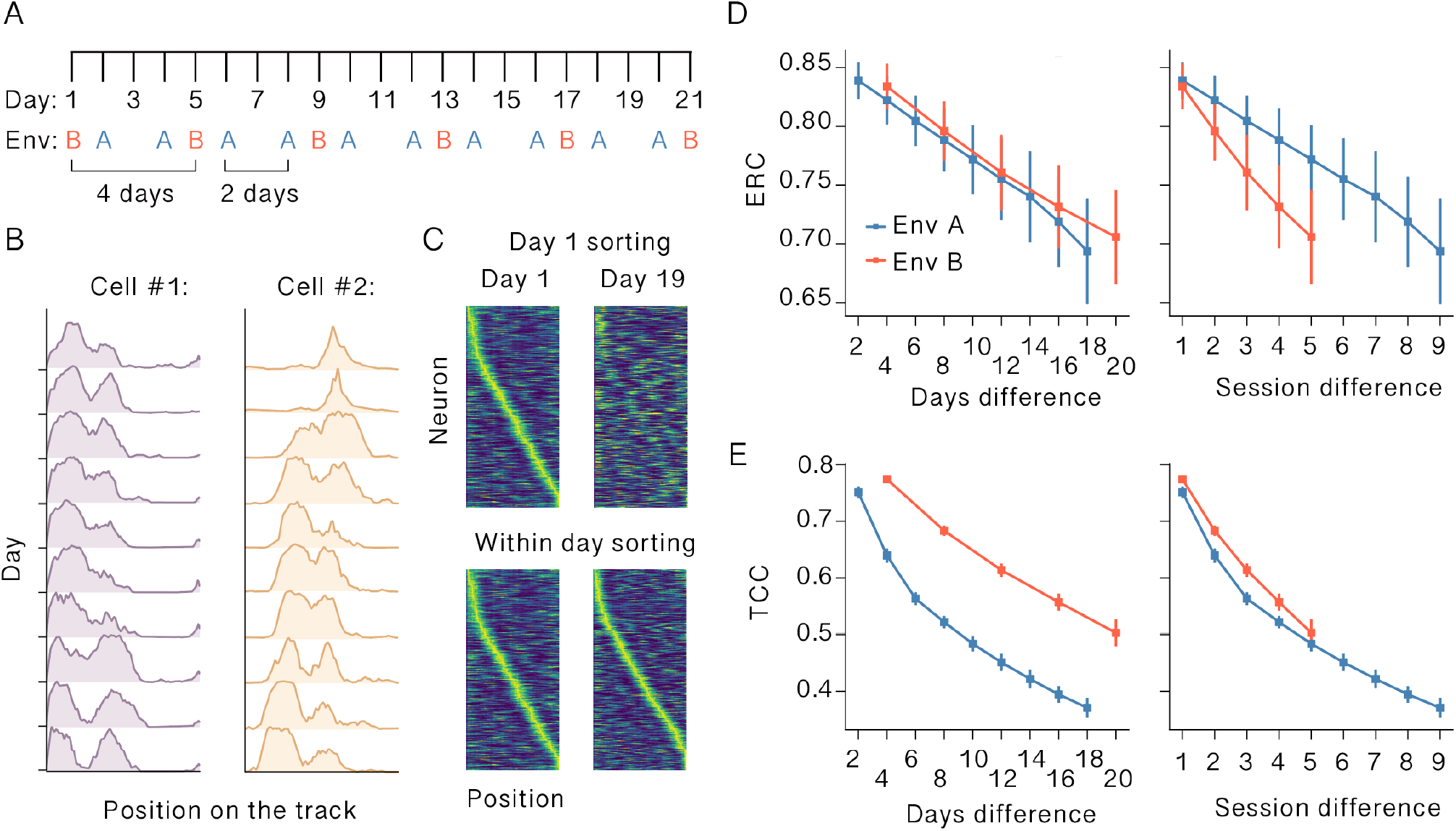
Predictive plasticity reproduces experience-dependent hippocampal representational drift. (A) Simulation schedule modeled after the longitudinal CA1 imaging paradigm of Geva et al. (2023). Env A, blue, was experienced every 2 days, whereas Env B, red, was experienced every 4 days. (B) Example tuning curves of two model neurons in Env A across recording days. Single-neuron spatial tuning changed gradually over repeated exposures. (C) Population tuning maps in Env A. Neurons were sorted either by their preferred position on Day 1 or by their preferred position within each day. Day 1 sorting revealed ordered place-field activity on Day 1, but not on Day 19. Sorting each day by its own preferred positions restored sequential population structure, indicating preserved spatial organization despite drift in individual tuning. (D) Ensemble rate correlations, ERC, plotted against elapsed days, left, or session difference, right. ERC decayed similarly across environments when measured by elapsed time, but differed when measured by intervening sessions. (E) Tuning curve correlations, TCC, plotted against elapsed days, left, or session difference, right. TCC differed across environments when measured by elapsed time, but became similar when measured by intervening sessions.

Despite this single-neuron drift, the population retained an ordered spatial representation. In Env A, sorting population activity by Day 1 preferred positions revealed disruption of sequential place-field organization, whereas sorting each day by its own preferred positions restored sequential population maps (Fig. 4C). Thus, as in the pattern-assembly task, spatial organization was preserved at the population level while the identity and preferred positions of individual neurons changed over time. Similar activity maps for both environments confirmed that spatial tuning drift occurred in Env A and Env B, while sequential population organization was preserved within each environment (Supplementary Fig. 2).

We then quantified representational stability using the same two measures used in the experimental study: ensemble rate correlation (ERC), and tuning curve correlation (TCC). ERC measures the similarity of each neuron’s mean firing rate across sessions, whereas TCC measures the similarity of each neuron’s spatial tuning profile across sessions. In the model, ERC decreased gradually as a function of elapsed time in both environments. When plotted against the number of separating days, Env A and Env B showed similar decay profiles. In contrast, when plotted against session difference, ERC decayed more rapidly in Env B than in Env A (Fig. 4D). This pattern is expected if ensemble-rate drift is primarily driven by elapsed time, because the same number of intervening sessions corresponds to a longer elapsed interval in the less frequently sampled environment.

Tuning curve stability showed the complementary pattern. TCC decreased gradually in both environments, but the decay differed between Env A and Env B when plotted against elapsed days. In contrast, when plotted against session difference, TCC decay became similar across the two environments (Fig. 4E). This indicates that tuning-curve drift depended more strongly on the amount of intervening experience than on elapsed time alone. Thus, the model reproduced the key dissociation observed in CA1: ensemble-rate stability was primarily time-dependent, whereas tuning-curve stability was more closely aligned with experience. This dissociation arises because mean firing rates are sensitive to cumulative plasticity-mediated fluctuations in synaptic drive, which are driven by ongoing stochastic spiking activity and can accumulate with elapsed time. In contrast, spatial tuning curves are reshaped primarily during active exposure to each environment, when position-specific inputs engage predictive E/I plasticity. We also found that the fraction of cells retaining their primary place field near the same position gradually decreased with increasing inter-session interval in both environments (Supplementary Fig. 3).

Together, these results show that predictive plasticity alone can reproduce key features of hippocampal representational drift across repeated environmental exposures. Single-neuron tuning changed gradually, but population-level spatial organization remained structured, and different stability measures showed separable dependencies on the exposure schedule.

### Coordinated drift is described by gradual rotation and translation in state space

The preceding analyses showed that the model reproduced gradual remodeling of spatial tuning across repeated exposures to the same environment. We next asked whether this drift reflected random degradation of the population code or a coordinated transformation of the underlying state space, inspired by recent analyses of CA1 recordings by Sylte et al. (2025). We visualized population-level changes by projecting activity from each simulated recording day onto the first three PCA axes (Fig. 5A). Later-day trajectories were progressively displaced and rotated relative to the Day 1 reference trajectory. However, after subspace alignment to the Day 1 reference, the trajectories from later days more closely matched the reference trajectory. Thus, although individual tuning curves changed over time, the population representation retained a substantial degree of geometric organization.

**Figure 5.**
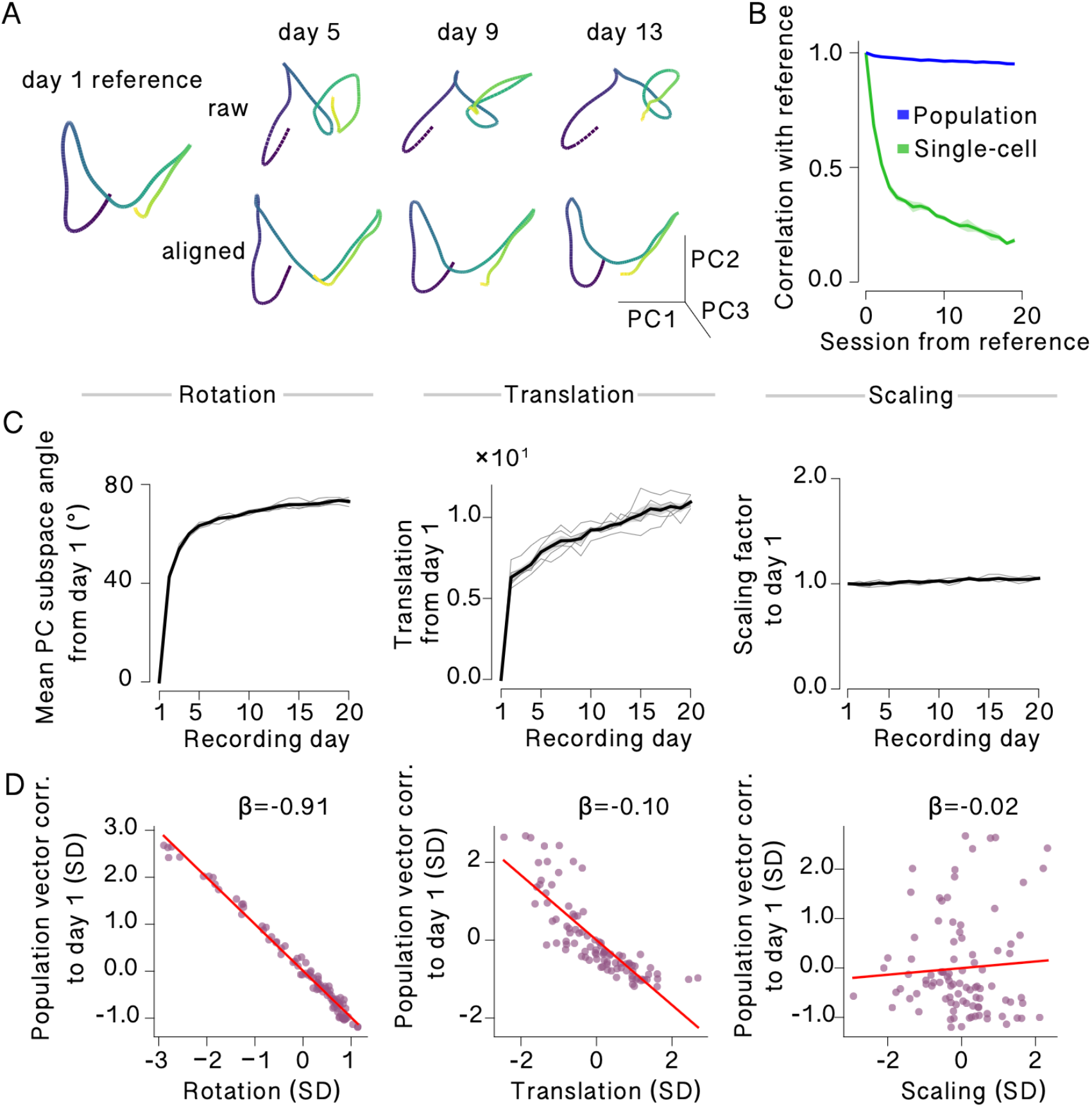
Coordinated drift preserves population geometry despite changes in single-cell tuning. (A) Population activity projected onto the first three PCA axes before and after subspace alignment, shown for example sessions. The left panel shows the Day 1 reference trajectory. For each later session, the upper row shows the raw trajectory and the lower row shows the aligned trajectory. (B) Correlation with the reference session for single-cell tuning and population geometry. Single-cell tuning correlation decreased more strongly over sessions than population geometry correlation, indicating that population-level structure was better preserved than individual tuning. (C) Similarity transformations relative to Day 1 across recording days. Rotation, quantified as the mean principal angle between PCA subspaces, and translation, quantified as centroid displacement, increased gradually over days, whereas uniform scaling remained approximately stable. Thin lines indicate individual simulations, and thick lines indicate the mean across simulations. (D) Linear mixed model relating population-vector correlation with the Day 1 reference to each similarity transformation while controlling for the other transformations. Rotation was the strongest predictor of the decline in population-vector correlation, with a weaker contribution from translation and little contribution from uniform scaling, similar to the experimental findings reported by Sylte et al. (2025). We performed 5 independent simulations.

We quantified this distinction by comparing single-cell tuning correlations and population geometry correlations with the reference session (Fig. 5B). Single-cell tuning correlation declined strongly over sessions, consistent with ongoing drift in individual spatial tuning. In contrast, population geometry correlation remained substantially higher across the same recording period. This dissociation indicates that the model did not simply lose spatial information over time. Rather, the identity and tuning of individual neurons changed while the structure of the population representation was partially preserved.

We then decomposed model drift into three similarity transformations: rotation, translation, and uniform scaling of the population state space. For each simulated recording day, we constructed a population activity matrix whose rows corresponded to spatial positions and whose columns corresponded to model neurons, and compared each day with Day 1. Rotation was quantified as the mean principal angle between the Day 1 PCA subspace and the corresponding subspace on each later day. Translation was quantified as the change of the population centroid relative to Day 1, and uniform scaling was quantified as the relative norm of mean-centered population activity. Rotation increased gradually across recording days, and translation also increased progressively over time (Fig. 5C). In contrast, scaling remained approximately stable, indicating that drift was not primarily caused by a global expansion or contraction of the population activity cloud.

Finally, we asked which of these transformations best explained the decline in population-vector correlation with Day 1. Using a linear mixed model that controlled for the other transformations, we found that rotation was the strongest predictor of population-vector drift, with a weaker contribution from translation and little contribution from uniform scaling (Fig. 5D).

Together, these results indicate that local predictive plasticity does not merely destabilize individual place fields. Instead, it produces a coordinated population-level drift in which the spatial code remains geometrically organized while the population state space gradually rotates and translates over time.

## Discussion

This work suggests that representational drift need not reflect random degradation of stored representations. Instead, it can emerge from the same plasticity mechanisms that maintain predictive E/I balance in the circuit. In the present learning rule, inhibitory synapses track total excitatory drive, implementing a form of homeostatic E/I co-tuning that has been established experimentally and theoretically (Vogels et al., 2011). As excitatory weights continue to change during ongoing learning, inhibitory plasticity gradually alters the relative net drive between competing representations, making individual neurons susceptible to reassignment. Changes in preference therefore occur not because excitation to the preferred pattern simply collapses, but because synaptic drive for that pattern is weakened as inhibition progressively strengthens onto the currently active assembly. At the population level, the same learning dynamics produced coordinated rotation and translation of state space, allowing population geometry to remain organized despite ongoing single-neuron drift.

This view differs from models in which drift is driven by externally imposed synaptic noise, stochastic weight perturbations, or Hebbian competition among fluctuating inputs (Kossio et al., 2021; Qin et al., 2023). In those frameworks, individual changes are effectively stochastic. Related work has shown that stable task information can be maintained despite unstable neural populations (Rule et al., 2020; Rule & O’Leary, 2022), and other models have linked assembly drift to STDP-based plasticity, dynamic attractor mechanisms, intrinsic excitability fluctuations, or synaptic volatility (Manz & Memmesheimer, 2023; Spalla et al., 2021; Mongillo et al., 2017; Delamare et al., 2024). Our results indicate that the CA1-like drift phenomenology does not require explicit synaptic rewiring or externally imposed weight perturbations. In the proposed model, predictive plasticity during active environmental exposure was sufficient to generate progressive changes in spatial tuning. This produced a day-dependent decline in ensemble rate correlation, while tuning-curve correlation was more closely aligned with the number of intervening environmental exposures.

More broadly, the framework suggests that different components of representational change can be dissociated because they are differentially shaped by experience-dependent synaptic learning and by the temporal structure of environmental exposure. In biological circuits, structural turnover may provide an additional source of long-timescale variability, and understanding how such processes interact with activity-dependent plasticity could help explain why population codes in different brain regions show different drift rates and different dependencies on behavioral state and experience. More generally, hippocampal place-code stability can be influenced by behavioral state and experience, as shown by exercise-dependent changes in place-code information and long-term stability (Rechavi et al., 2022; de Snoo et al., 2023).

The model makes several experimentally testable predictions. First, neurons with weaker current response selectivity should be more likely to change their preferred representation on subsequent exposures, because low selectivity reflects reduced net drive for the current preferred pattern. This prediction can be tested in longitudinal recordings by asking whether single-cell selectivity in one session predicts remapping probability in the next. Second, partial suppression of inhibitory plasticity should slow drift rates while preserving pattern selectivity, because inhibitory weight changes are the proximal driver of the excitatory learning-signal reversal that underlies preference change. Finally, although structural synaptic turnover was not required in the present model, it may contribute to long-timescale drift in biological circuits.

Together, these findings suggest that representational drift reflects an intrinsic consequence of predictive E/I plasticity rather than a breakdown of memory stability. By continually rebalancing excitation and inhibition, the same local learning rule can preserve structured population codes and organized state-space geometry while gradually reassigning the representational roles of individual neurons.

## Methods

### Neural network model

Throughout this study, we used recurrent neural networks consisting of N = 400 Poisson neurons. Each neuron generated spikes according to a non-stationary Poisson process with rate *φ*(*u*), where *φ*(•) is a sigmoidal function. The membrane potential *u* of neuron *i* at time *t* is given as follows:

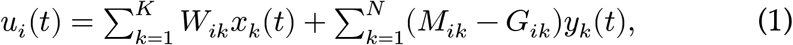

where *K* is the number of input neurons. Three matrices **W** ∈ ℝ^*N*×*K*^, **M** ∈ ℝ^*N*×*N*^, and **G** ∈ ℝ^*N*×*N*^ represent the weights of excitatory afferent synaptic connections, excitatory recurrent synaptic connections and inhibitory recurrent connections, respectively, on neurons in the recurrent network. Initial values of **W**, **M**, and **G** were sampled from Gaussian distributions with mean zero and standard deviations 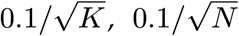, and 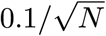, respectively, and were then rectified by taking their absolute values. These synaptic connections are all-to-all. In terms of the kernel function

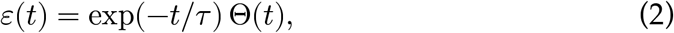

recurrent input and afferent input to neuron *i* are calculated as

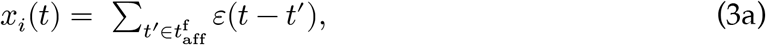

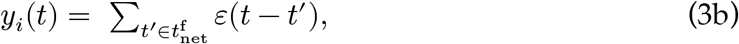

where *τ* stands for the membrane time constant, 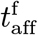 and 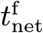 for the time sets of afferent and recurrent presynaptic spikes, and Θ(•) for the Heaviside function. Throughout this study, *τ* = 15 ms.

The instantaneous firing rate *fi*(*t*) of each neuron is given as

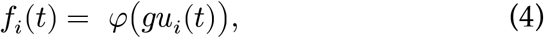

in terms of sigmoidal response function *φ*:

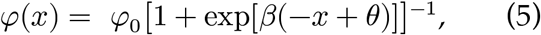

with the maximum instantaneous firing rate *φ*_0_ of 50 Hz. The values of constant parameters were set as *g* = 3, *β* = 5, and *θ* = 1.

### Synaptic plasticity rules

To predict the firing rate of the postsynaptic neuron, the different types of synapses obey similar learning rules in the present network model. Given the postsynaptic potentials as

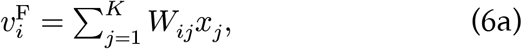

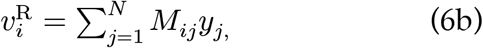

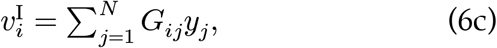

the weights of the corresponding synapses are modified according to the following equations:

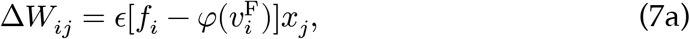

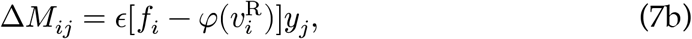

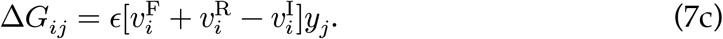

The learning rate was set to *ϵ* = 10^−4^ in Figs. 2 and 3, and *ϵ* = 3 × 10^−4^ in Figs. 4 and 5.

### Details of patterned input simulations

For Figures 2 and 3, we used three discrete input patterns, each lasting 500 ms. Each pattern activated a distinct, non-overlapping subset of 100 input neurons at a fixed rate of 30 Hz, while all remaining input neurons were silent. To visualize population-level representations, neurons were sorted by their preferred pattern, defined as the pattern whose presentation evoked the highest time-averaged correlation with the neuron’s activity trace. Assembly overlap was computed for each reference assembly as the ratio of the mean firing rate of reference assembly members at a given snapshot to that at the reference snapshot, averaged over members whose selectivity exceeded the mean selectivity of that assembly at the reference snapshot.

### Trial-resolved drive decomposition and preference tracking

In Figure 3, we recorded trial-averaged firing responses, excitatory drive, inhibitory drive, net E/I drive, and excitatory prediction errors for each neuron during the drift phase. Pattern-conditioned values were estimated with a sliding-window average over the five most recent presentations of each pattern, and the preferred pattern was defined as the pattern evoking the largest mean firing response. Stable changes were defined as changes in preferred pattern that persisted for at least two consecutive trials before and after the transition. We then computed transition-triggered averages around detected changes and asked whether current selectivity predicted change on the following trial. Selectivity was defined as the difference between the largest and second-largest pattern-conditioned values for response, excitatory drive, or inhibitory drive, and change probability was computed after binning trials by each selectivity measure.

### Details of simulating place field sequences

In Figure 4, we used a spatial variant of the same model to reproduce the experimental paradigm of Geva et al. (2023). Afferent input was provided by 800 spatially tuned input neurons with Gaussian tuning curves tiling a virtual linear track of 400 positions. Two environments were encoded by non-overlapping subsets of input neurons: neurons 1–400 for Env A and neurons 401–800 for Env B. The protocol followed the 21-day schedule of Geva et al. (2023): Env A was visited every 2 days, whereas Env B was visited every 4 days. Each session consisted of a training epoch with learning enabled, followed by a frozen-weight test epoch during which spatial tuning curves were measured.

Representational stability was quantified using the same two metrics as Geva et al. (2023). Ensemble rate correlation (ERC) was computed as the Pearson correlation between vectors of mean firing rates across neurons, whereas tuning curve correlation (TCC) was computed as the mean across neurons of the Pearson correlation between spatial tuning curves. Both metrics were evaluated as a function of elapsed calendar days and intervening sessions, separately for each environment. Sorted tuning maps were generated by normalizing each neuron’s tuning curve within session and sorting neurons by their preferred track position.

### Population geometry and similarity transformation analyses

In Figure 5, to quantify coordinated drift of the spatial population code, for each session, we constructed a position-by-neuron population activity matrix after normalizing each neuron’s tuning curve within session. Single-cell tuning correlation was computed as the mean correlation between each neuron’s tuning curve in a given session and on Day 1. Population geometry correlation was computed by comparing representational dissimilarity matrices across spatial positions with the Day 1 reference.

For visualization, Day 1 activity was used to define a three-dimensional PCA space, and activity from later sessions was projected into this space before and after alignment to the Day 1 reference trajectory. To quantify coordinated drift, we decomposed changes relative to Day 1 into rotation, translation, and uniform scaling. Rotation was measured as the mean principal angle between PCA subspaces, translation as the displacement of the population centroid, and scaling as the relative norm of mean-centered population activity. Finally, we used a linear mixed model to estimate how each transformation contributed to population-vector drift while controlling for the others, with simulation identity included as a grouping factor.

### Simulation details

All simulations were performed in customized Python3 code written by TA with numpy 2.3.5 and scipy 1.17.0. Differential equations were numerically integrated using an Euler method with integration time steps of 1 ms.

## Acknowledgments

The authors thank Carl E. Schoonover and Andrew J. P. Fink for their valuable discussions. This work was supported by Wellcome Trust 200790/Z/16/Z, EPSRC EP/R035806/1, ERC MotorAdapt 101169605 to C.C.

## Author Contributions

T.A. and C.C. conceived the study and wrote the paper. T.A. performed the simulations and data analyses.

## Supplementary Figures

**Supplementary Figure 1:**
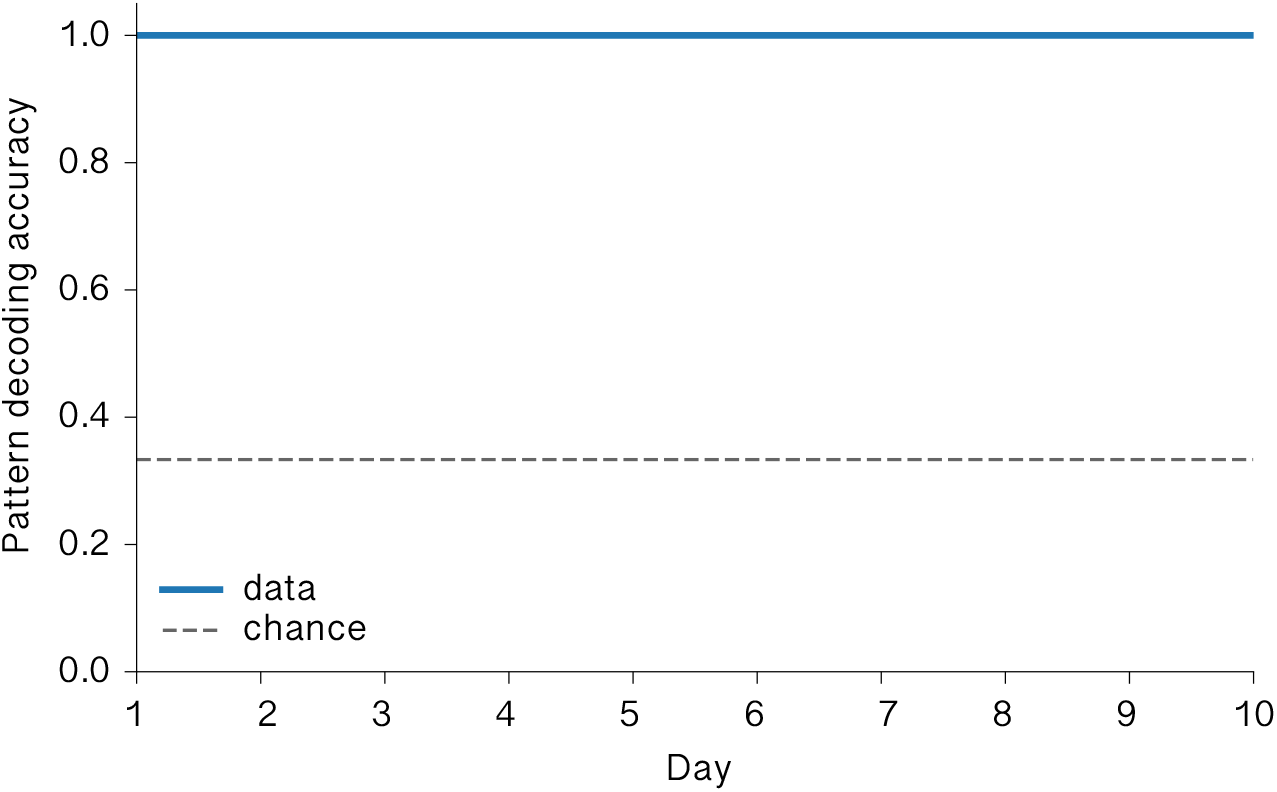
Pattern identity remains decodable throughout assembly drift. Pattern decoding accuracy across the drift period in the patterned-input simulation. Decoding was performed from population activity on each day. The gray dashed line indicates chance-level performance.

**Supplementary Figure 2:**
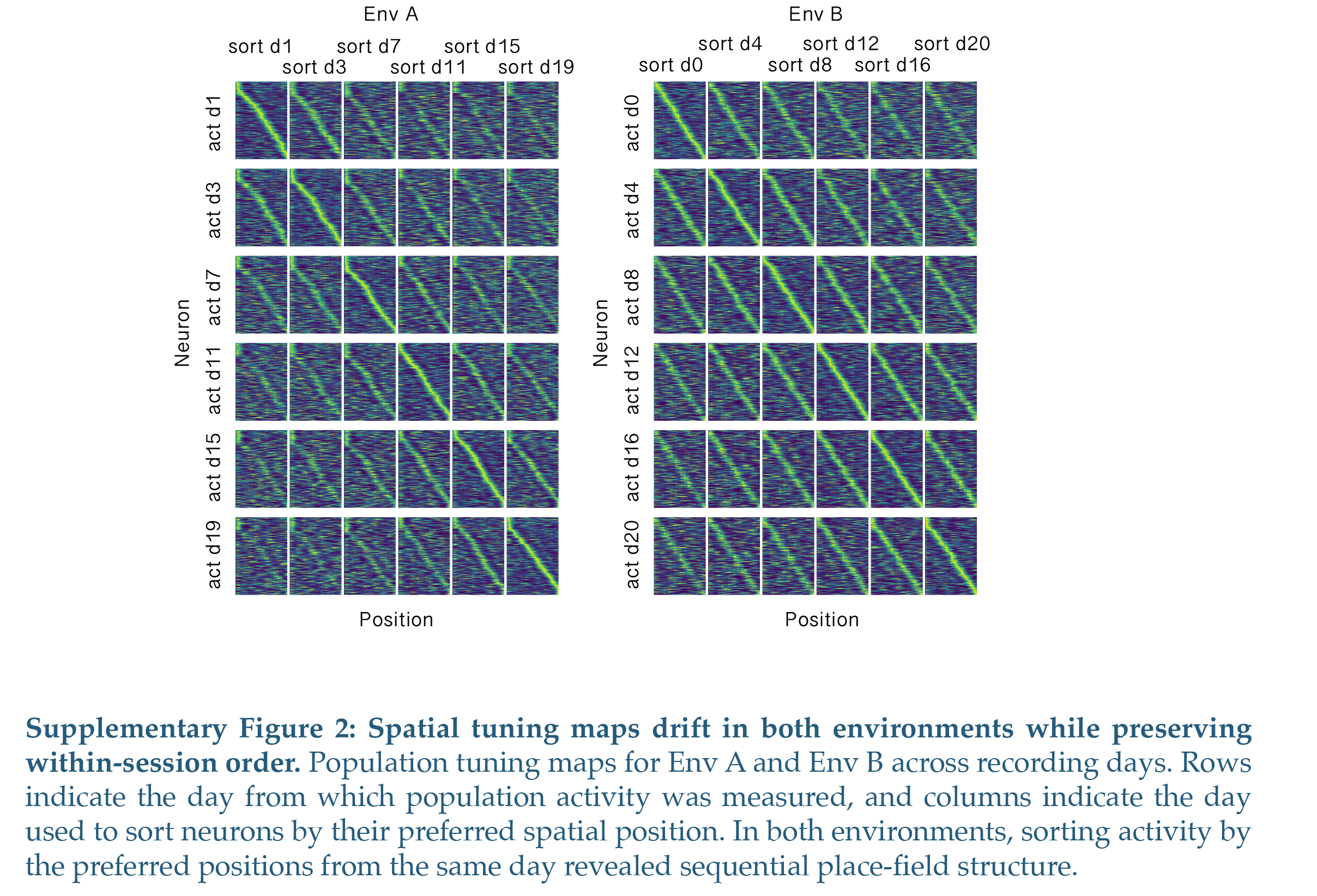
Spatial tuning maps drift in both environments while preserving within-session order. Population tuning maps for Env A and Env B across recording days. Rows indicate the day from which population activity was measured, and columns indicate the day used to sort neurons by their preferred spatial position. In both environments, sorting activity by the preferred positions from the same day revealed sequential place-field structure.

**Supplementary Figure 3:**
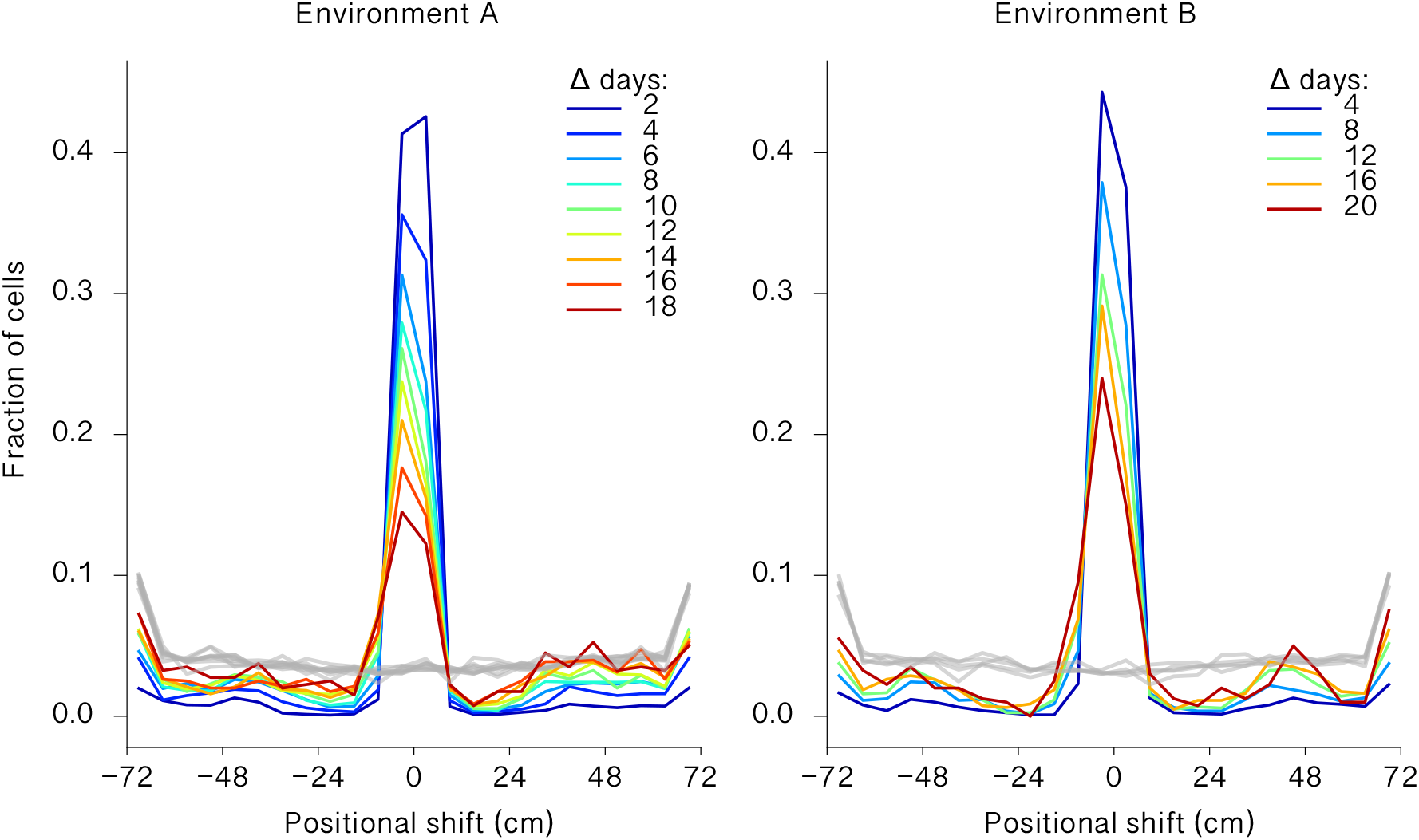
Place-field positional shifts increase with inter-session interval. Distribution of primary place-field positional shifts between pairs of sessions in Env A and Env B are shown. Colored lines indicate session pairs separated by different elapsed time intervals. In both environments, the distribution of positional shifts broadened as the interval between sessions increased. Gray lines indicate shuffled controls in which cell identities were independently permuted within each day.

